# Estimating heterogeneous treatment effects by balancing heterogeneity and fitness

**DOI:** 10.1101/333278

**Authors:** Weijia Zhang, Thuc Le, Lin Liu, Jiuyong Li

**Affiliations:** School of Information Technology and Mathematical Sciences, University of South Australia, Adelaide, Australia; Centre for Cancer Biology, University of South Australia, Adelaide, Australia

## Abstract

Estimating heterogeneous treatment effects is an important problem in many medical and biological applications since treatments may have different effects on the prognoses of different patients. Recently, several recursive partitioning methods have been proposed to identify the subgroups that with different responds to a treatment, and they rely on a fitness criterion to minimize the error between the estimated treatment effects and the unobservable true effects. In this paper, we propose that a heterogeneity criterion, which maximizes the differences of treatment effects among the subgroups, also needs to be considered. Moreover, we show that better performances can be achieved when the fitness and the heterogeneous criteria are considered simultaneously. Selecting the optimal splitting points then becomes a multi-objective problem; however, a solution that achieves optimal in both aspects are often not available. To solve this problem, we propose a multi-objective splitting procedure to balance both criteria. The proposed procedure is computationally efficient and fits naturally into the existing recursive partitioning framework. Experimental results show that the proposed multi-objective approach performs consistently better than existing ones.

**Author summary:** The effects of a treatment are often not the same for different individuals with different gene expressions. Learning to predict the heterogeneous treatment effects from clinical and expression data is an important step towards personalized medical treatment. Existing computational methods are not ideal for the task because they do not address the interpretability of the model and do not consider the limited sample sizes in biological and medical applications. Our method addresses these issues and achieves superior performance in analyzing the treatment effects of radiotherapy on breast cancer patients.

## Introduction

Treatment effect estimation is a fundamental problem in scientific research. Biologists use it to study the regulatory relationships between numerous genes, and medical researchers rely on it to determine whether a treatment is effective for the patients [1].

Traditionally, treatment effects are averaged over the entire population [2]. However, understanding their heterogeneities is often beneficial because different individuals may react differently towards a particular treatment. For example, although radiotherapy is an effective treatment for most breast cancer patients, many patients do no benefit from the treatment because of their different gene expression patterns [3]. Understanding the heterogeneities in the treatment effects of radiotherapy could prevent patients from suffering unnecessary side effects.

It is desirable to apply principled data mining algorithms to identify treatment effect heterogeneities [4]. Tree-based recursive partitioning methods [5], originally proposed for regression and classification tasks, are perfect candidates for modeling heterogeneous treatment effects. Unlike methods which have strong predictive power but are difficult to interpret, tree-based methods often excel on both aspects. Their model output, trees, can be easily interpreted by human experts, which is important in many applications that require treatment effect estimation, such as biomedical and social studies.

Existing recursive partitioning methods cannot be applied to treatment effect estimation directly. When building regression and classification trees, algorithms have access to the target variables in the training data. Unfortunately, such information is not available when estimating treatment effects because an individual can either be treated or not treated, but not both at the same time. In other words, only one of the two potential outcomes is observable but both outcomes are needed for the estimation [6].

Following the established paradigm of maximizing the fitness of the tree model at each splitting point, a number of recursive partitioning methods have been proposed to estimate heterogeneous treatment effects by utilizing an alternative fitness criterion [7, 8]. Specifically, these methods rely on a surrogate function to minimize the error between the estimated treatment effects and the unobservable true effects. The surrogate loss function approximates the true error when the available sample size is sufficiently large. However, the available sample size decreases rapidly as the tree grows and the surrogate function may not accurately estimate the model error.

Understanding the heterogeneity in treatment effects has important real-world implications. For example, consider two models regarding the treatment effects of radiotherapy on cancer patients (Fig. 1). The first model divides the patients according to the expression level of *gene*_1_, and identifies two sub-populations with similar treatment effects. The second model places the split at *gene*_2_, and discovers subgroups with significantly different treatment effects. If the errors of both models are acceptable, the second model should be preferred because it provides more insight into how treatment effects vary among different subgroups.

**Fig 1.**
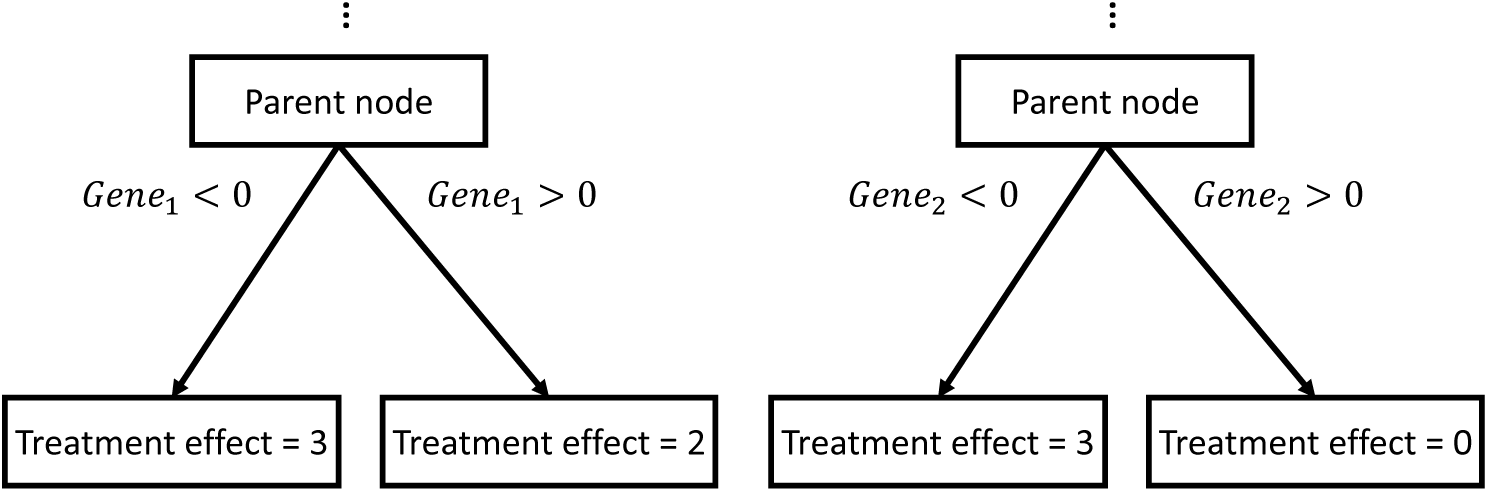
Illustration of two models for estimating heterogeneous treatment effect. The first model preferred fitness while the second one prioritized heterogeneity.

Unfortunately, algorithms relying on the fitness criterion often prefer the first model to the second one (Fig. 2). When choosing a splitting point, fitness based methods would prefer the first model because splitting at *gene*_1_ achieves lower estimation error than splitting at *gene*_2_. However, it is important to note that the estimated errors may not truthfully reflects the true errors of the models since the surrogate function is only accurate when the sample sizes are sufficiently large but the available sample sizes decreases quickly as the trees grow.

**Fig 2.**
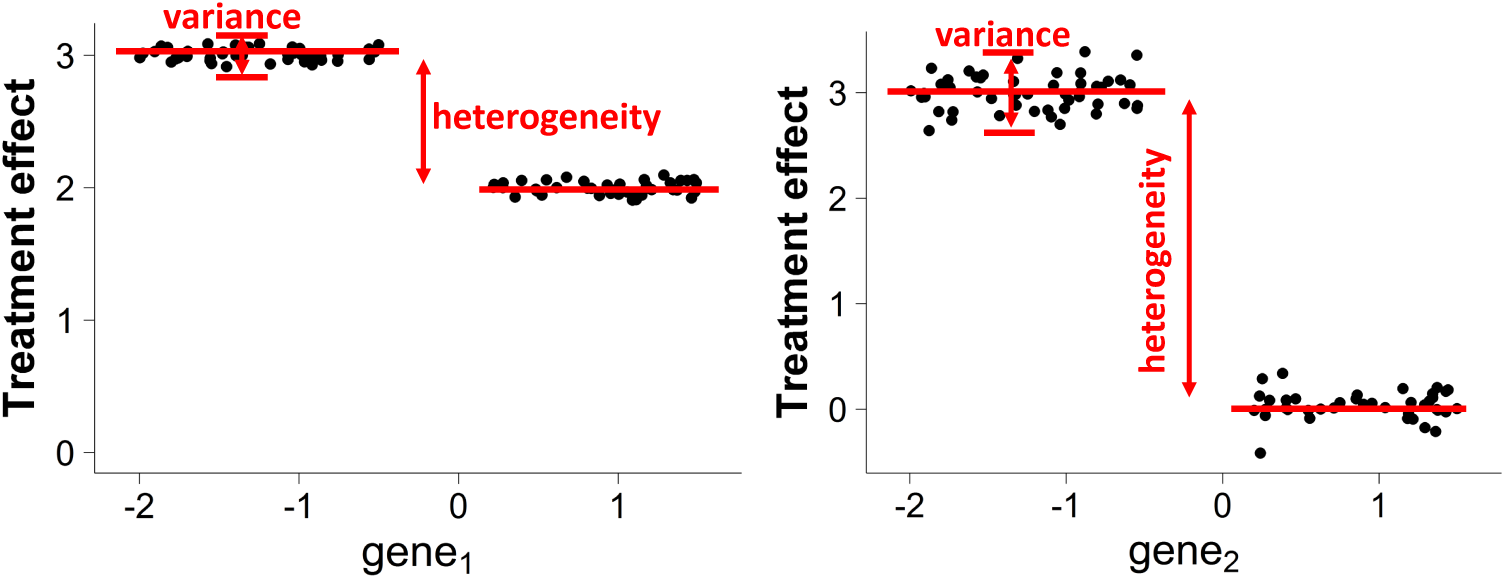
Estimated treatment effects for models in Fig. 1. When the sub-populations are split at *gene*2, the estimated error is slightly larger than splitting at *gene*_1_. This explains why existing methods prefer Model 1. However, the heterogeneity of treatment effects is ignored in this criterion.

The heterogeneity of treatment effects needs to be explicitly considered in recursive partitioning. A method would prefer the second model to the first one if it directly maximizes the heterogeneities, i.e., maximizing the difference between the treatment effects of two child nodes. Thus, utilizing the heterogeneities will often produce a more informative model than relying on the fitnesses.

Heterogeneity and fitness need to be simultaneously considered in order to generate a informative and reliable model. If an algorithm relies exclusively on maximizing the heterogeneity, it is likely to favor models with spuriously high treatment effect differences and unacceptable model errors. In the previous example, there may exist another variable which leads to unacceptable high error, but also results in more significant heterogeneities because of noises and outliers.

Finding the optimal splits then becomes a multiple-objective problem: the first objective is to maximize the fitness of the model and the second objective is to maximize the heterogeneity among the subgroups. Unfortunately, in real-world applications, splitting points that maximize both objectives simultaneously are often not achievable.

In this paper, we first propose the Maximizing Heterogeneity (MH) criterion which aims to maximize the differences in treatment effects. Then we propose the multi-objective (MO) criterion which considers both the heterogeneities and the fitnesses during the partitioning procedure. When solutions which maximize heterogeneity and fitness simultaneously are not achievable, MO chooses splitting points that achieve high scores in both criteria by allowing a degree of tolerance in their dominance relationships.

The proposed criteria are compared with existing ones using both simulated and real-world datasets. Experiment results demonstrate that while MH performs better than existing ones in many cases, it is prone to error when the differences in treatment effects become small. With fitness and heterogeneity are both taken into consideration, MO performs consistently better than compared methods.

## Methods

### Preliminaries

In this section, we introduce necessary definitions and results for heterogeneous treatment effect estimation.

Let *W*_*i*_ ∈ {0, 1} denote the treatment assignment, *Y*_*i*_ denote the observed outcome, and **x**_*i*_ = {*x*_*i*1_, *…, x*_*ip*_} denote the pre-treatment covariates. The data consists of i.i.d. samples (*Y*_*i*_, *W*_*i*_, **x**_*i*_), for *i* = 1, *…, N*. For the sake of simplicity, the subscript *i* will be omitted when the context is clear.

Let *Y* ^(*W*)^ denote the potential outcome if an individual has received the treatment *W*, then the observed outcome *Y* can be described as *Y* = *WY* ^(1)^ + (1 – *W*)*Y* ^(0)^. Although each sample is associated with two potential outcomes *Y* ^(1)^ and *Y* ^(0)^, only one of them can be realized as the observed outcome *Y*.

The average treatment effect (ATE) is defined as the expected outcome if the entire population were treated minus the outcome if they were not treated [6]:

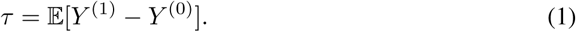

Since only one of the two potential outcomes can be observed, Equation 1 is *counterfactual* and cannot be estimated straightforwardly. When the treatment assignment is completely random, i.e., (*Y* ^(0)^, *Y* ^(1)^) ╨ *W*), the average treatment effect can be estimated as *τ* = 𝔼 (*Y |W* = 1) – 𝔼 (*Y |W* = 0).

However, the treatment assignment is often not randomized. In such cases, the unconfoundedness assumption [6] is needed in order to estimate treatment effect in these circumstances:

#### Assumption 1

*W* ╨ (*Y* ^(0)^, *Y* ^(1)^)|**x**.

With the assumption, an unbiased ATE estimation can be achieved with the help of propensity score [9]. The propensity score is defined as *e*(**x**) = *Pr*(*W* = 1|**x**), the probability of treatment assignment conditioning on the covariates.

The propensity score can then be estimated with a variety of methods. Some popular choices include logistic regression, random forests, and boosting [10].

When treatment effects are heterogeneous across the population, estimating the conditional average treatment effect (CATE) [6] in various subpopulations defined by the possible values of the covariates **x** often provides more insight than estimating the ATE on the entire population. Specifically, CATE is defined as:

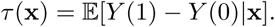

Recursive partitioning provides an ideal way for estimating CATE. Starting from the root node containing the entire population, a tree model is constructed by recursively splitting the node into two disjoint child nodes. By the end of the procedure, the subpopulations with heterogeneous treatment effects are naturally presented in the leaves of the model. For each leaf node, *τ* (**x**) can be estimated by calculating the ATE using only the samples within the node as follows:

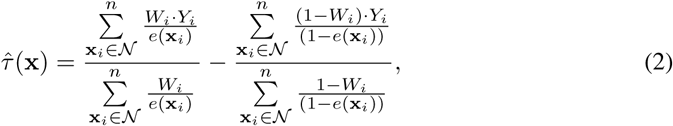

where the treatment propensity *e*(**x**_*i*_) is either known from experimental design or estimated from observational data.

The core component of a recursive partitioning model is the splitting criterion. At each split, the splitting criterion relies on a scoring function to evaluate the qualities of all potential splitting points. The recursive partitioning model then makes the split at the splitting point with the highest score.

The fitness criterion, one of the most widely adopted splitting criteria, aims to maximize the fitness of the model by minimizing the mean squared error(MSE). However, since the true treatment effects are not observable, the MSE cannot be estimated straightforwardly. In [**?**, 7], the authors observed that under Assumption 1, it can be obtained that

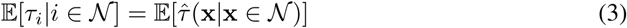

Relying on Equation 3, [7] proposed to utilize an alternative scoring function to estimate the error as:

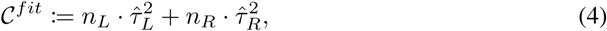

where *τ*_*L*_ and *τ*_*R*_ are the estimated treatment effects, *n*_*L*_ and *n*_*R*_ are the numbers of samples in the left and right child node.

### Proposed method

The problem of the fitness criterion 𝒞^*fit*^ is that Equation 3 is only valid when the sample size is sufficiently large. Unfortunately, in recursive partitioning the available sample size decreases quickly as the tree grows. The problem is even more severe in many applications where treatment effect estimation are required such as biological and sociology studies, where the sample sizes are already small to start with.

Inspired by the observations in the first section, we propose that a heterogeneity criterion which maximizes the differences in treatment effects of the child nodes, should be explicitly considered during the recursive partitioning process. Specifically, the heterogeneity criterion favors the split leading to the largest differences between the treatment effects of the two child nodes:

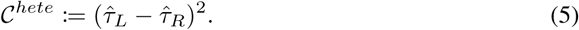

In the following sections, we will refer the criterion in Equation 5 as Maximizing Heterogeneity (MH).

MH is adept at identifying significant differences of treatment effects among sub-populations because it explicitly rewards splitting points with large heterogeneities. As will be demonstrated in the next section, using the MH criterion in recursive partitioning achieves better performances than using the fitness criterion when the underlying treatment effect heterogeneities are large, especially during the first few splits of the trees. Unfortunately, the performance of MH may decrease when the underlying heterogeneities are small.

As the tree grows and the heterogeneities become smaller, MH is prone to be misled by spurious heterogeneities. This is because MH does not place any consideration on the fitness of the model. In other words, the MH criterion would select a splitting point with high heterogeneity even if it also has high mean squared error. Consider the example of Fig. 2, if there exists another covariate *gene*_3_ with spuriously high heterogeneity, MH will use it to split and produce an unreliable model with high deviation from the truth.

The above problem could be avoided if the heterogeneity and the fitness are taken into consideration simultaneously. As 𝒞^*hete*^ identifies a splitting point with high heterogeneity, 𝒞^*fit*^ can be used as a quality assurance to double-check whether the splitting point also increases the fitness of the model. Therefore, an ideal splitting point should achieve the highest quality in terms of both the fitness criterion and the heterogeneity criterion.

However, the ideal splitting point is often not achievable in real-world data. Because the fitness criterion relies on the surrogate function, it may not be accurate when the sample size is not sufficiently large. Moreover, the heterogeneity criterion is vulnerable to noises and outliers in the data.

To solve this problem, we propose the Multi-Objective (MO) splitting criterion. Specifically, when a splitting point that maximizes both the fitness and the heterogeneity criteria is not available, MO aims to find a good compromise that has high scores in both aspects.

Let 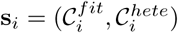 denote a fitness and heterogeneity scores for the *i*-th splitting point, and let *𝒮* be the set containing the score pairs corresponding to all potential splitting points. The goal is to find a 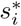 from 𝒮, where both the fitness and heterogeneity of 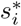 are as close to their maximum possible values as possible. To achieve this, we propose to utilize the ∊-dominance relationship [11].

#### Definition 1

(*ε*-dominance). *Score pair s*_*i*_ ∈ *𝒮 is said to* ∊*-dominate s*_*j*_ ∈ *𝒮 for some* ∊ = (∊_1_, ∊_2_), *denoted as s*_*i*_ *>*_∊_ *s*_*j*_, *if and only if:*

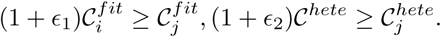

The *ε*-dominance relationship provides algorithms the abilities to control the amount of “tolerance” in the ordering of score pairs. Fig. 1 illustrates the comparison of *ε*-dominance with Pareto-dominance [12]. Intuitively, in order for score pair *s*_*i*_ to *ε*-dominate *s*_*j*_, both of *s*_*i*_’s components must be at least larger than those of *s*_*j*_’s by a margin specified by *ε*. Therefore, by specifying suitable *∊* values, MO is more resistant to the noises and outliers in the data.

With Definition 1, the *ε*-optimal set *𝒮*^*\**^ of *𝒮* is a set where all elements in *S* is *ε*-dominated by at least one element of *𝒮* ^*\**^, and all elements in *𝒮* ^*\**^ are in the Pareto-set of *𝒮*:

#### Definition 2

(*ε*-optimal set). *Let S* ⊆ ℝ^2^ *be a set of score vectors. Then the E-splitting set S*^*\**^ *is defined as follows:*

1. *Any score s* ∈ *S is* ∊*-dominated by at least one score s*^*\**^ ∈ *S*^*\**^, *i.e.*

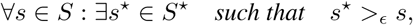
2. *Every score s*^*\**^ ∈ *S*^*\**^ *are not Pareto-dominated by any score s* ∈ *S, i.e.*

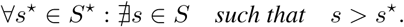

Comparison of Pareto-set and *ε*-optimal set are illustrated in Fig. 4, the top left panel depicts the elements in *𝒮* and its corresponding Pareto-set, and other panels describe the *ε*-optimal set with various *ε*. Compared to the Pareto-set, *ε*-optimal set contains significantly smaller number of elements. When *ε* is sufficiently small, the *ε*-optimal set is equivalent to the Pareto-optimal set [12].

**Fig 3.**
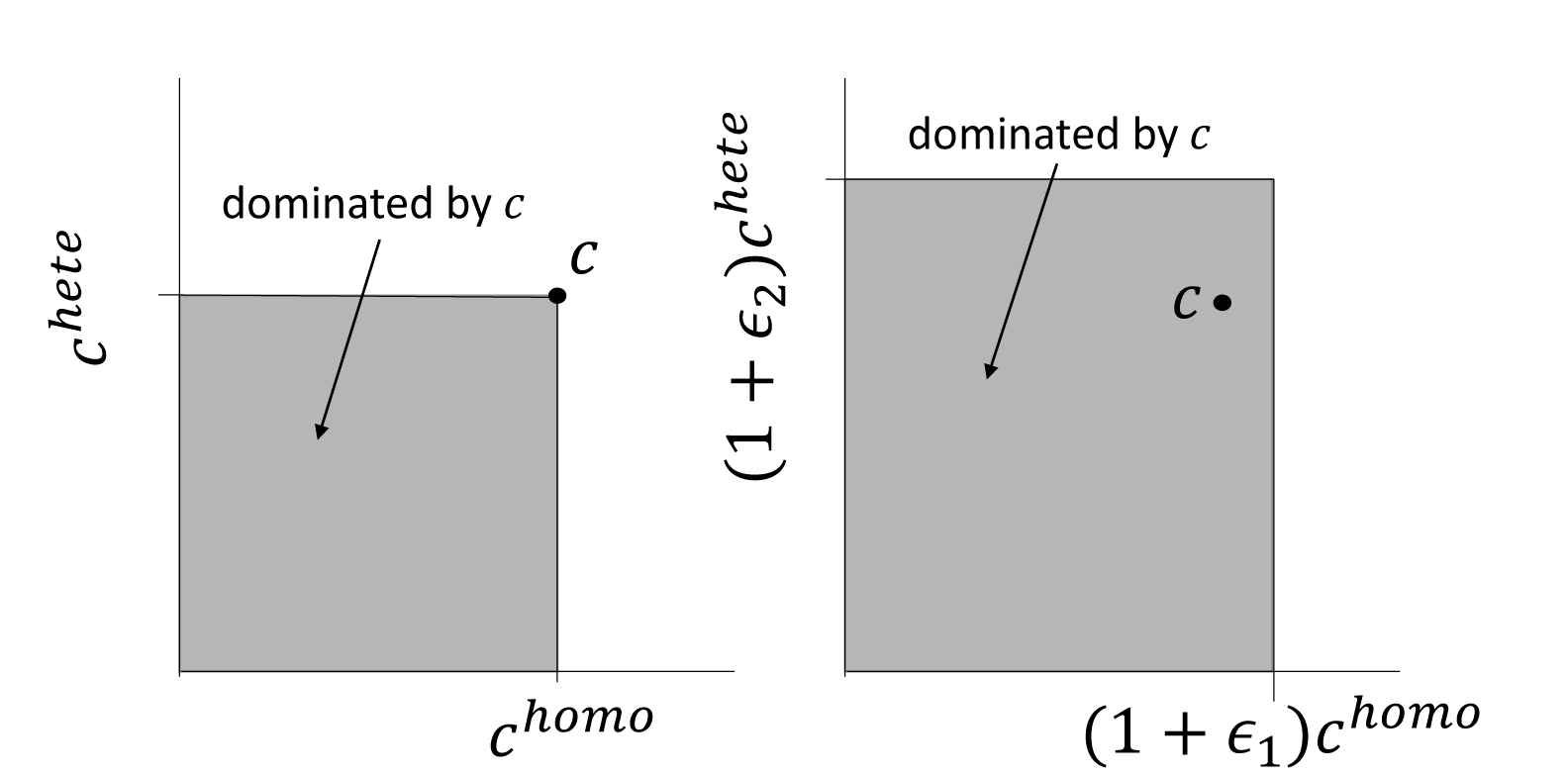
Comparison of Pareto-dominance and *ϵ*-dominance.

**Fig 4.**
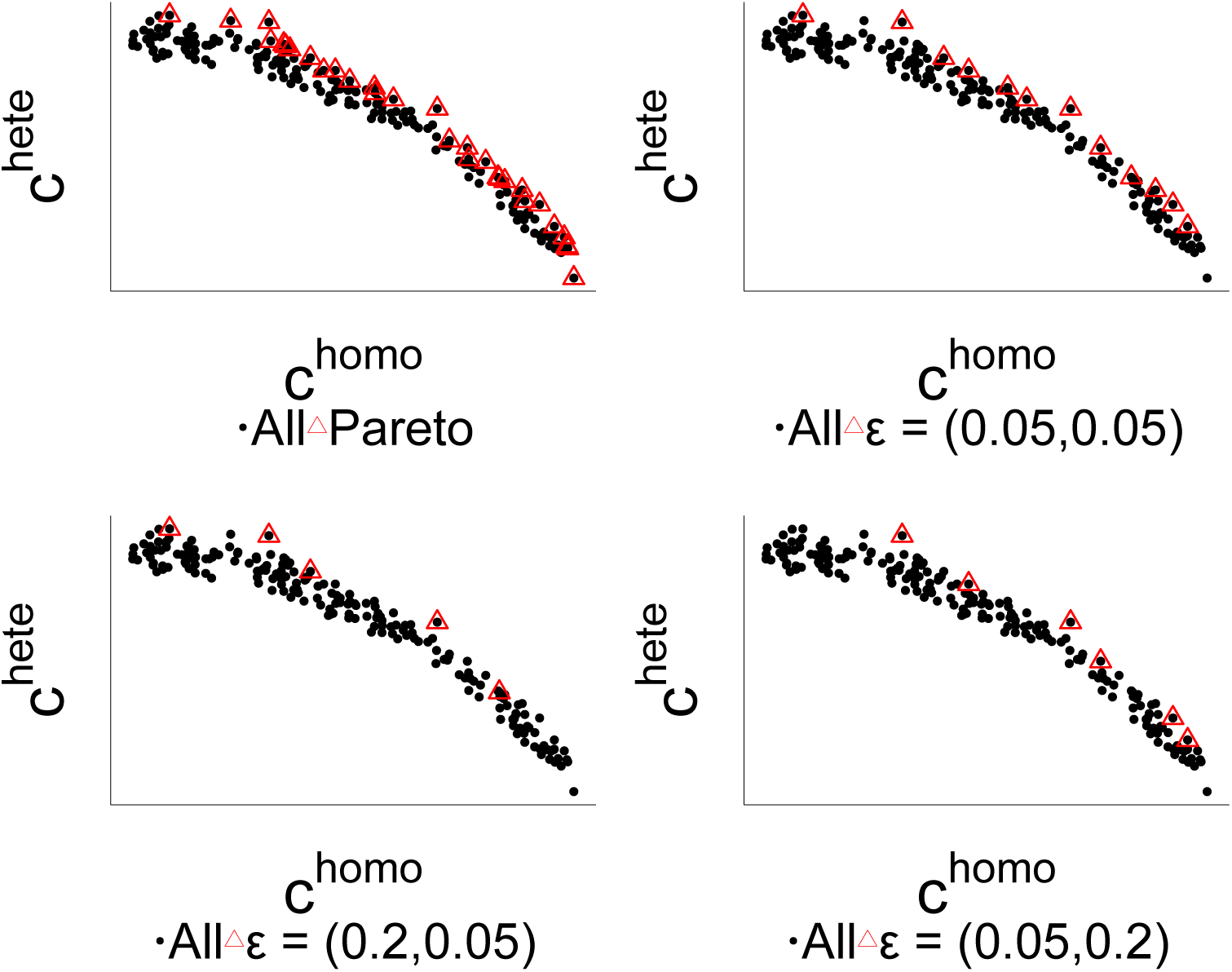
Comparison of Pareto-set and the *ϵ*-optimal set with various *ε* values.

We now discuss how to maintain the *ε*-optimal set while scanning through all the potential splitting points. This can be achieved by dividing the two dimension search space into squares of size 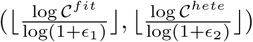, and only keeps one element which are not *ε*-dominated by others within the squares. We present the details in Algorithm 2.

Algorithm 2 has two important properties. Firstly it is guaranteed to converge to the *ε*-optimal set. Secondly, it is guaranteed that at all times the algorithm only needs to deal with a small number of score pairs.

#### Theorem 1.

*Let S be the set of all score pairs for all possible splitting points. Then the output of Algorithm 2 S*^*\**^ *is an ε-optimal set of S with bounded size:*

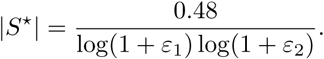

**Proof Sketch.** *On the coarse level, the search space is discretized into two-dimensional squares of size* 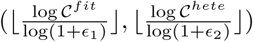, *where each vector uniquely belongs to one of the squares. Applying the ε dominance relation on these spaces, the algorithm always maintains a set of non-dominated squares, thus guaranteeing the ε-optimal property. On the fine level at most one element is kept as a representative vector in each square. Within a square, the representative vector can only be replaced another one if it is ε-dominated, thus guaranteeing convergence.*

The theorem indicates that the size of set *S*^*\**^ is small and irrelevant of the total number of score pairs. For example, if ∈_1_ and ∈_2_ are set to 0.2, then *S*^*\**^ is guaranteed to contain less than a hundred score pairs. Since Algorithm 2 needs to be executed at each split of the tree, its time complexity is crucial to the computational efficiency of the MO. Thanks to Theorem 1, the time complexity of MO is comparable to existing criteria since it only introduces small amount of extra computation cost for maintaining the *ε*-optimal set.

#### Algorithm 1: Tree construction procedure

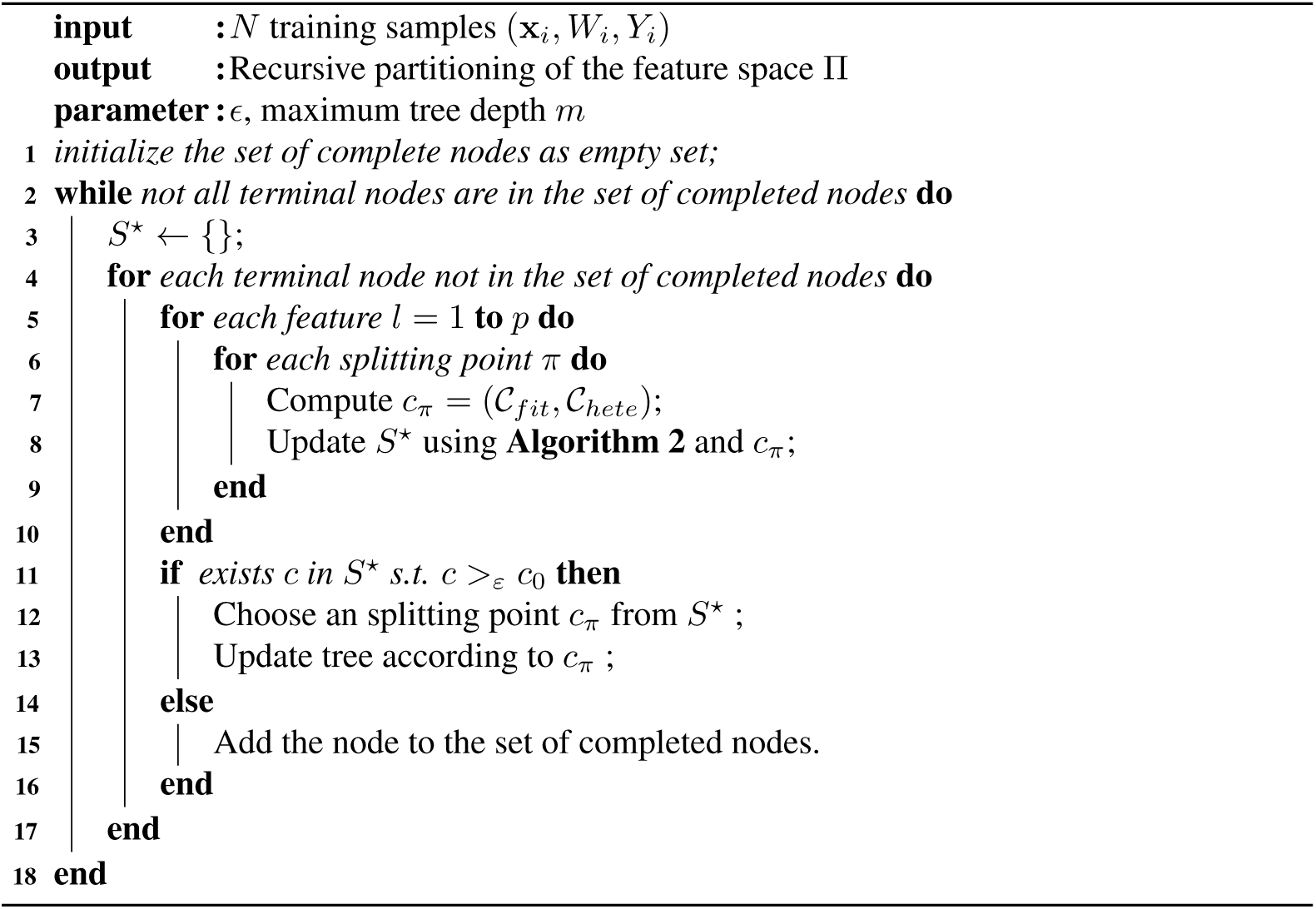

Although the *ε*-optimal set is guaranteed to be of small size, it is still necessary to select one splitting point from *S*^*\**^. To emphasize the utility of the heterogeneity criterion, in our implementation splitting points with the highest 𝒞^*hete*^ scores are chosen from the set. In the next section, it can be seen that this simple strategy already achieves superior performance; however, the results also indicates that further improvement can be achieved with more sophisticated strategies.

Finally, we summarize the multi-objective tree construction procedure in Algorithm 1. The structure of splitting procedure remains similar to the CART [5] method. However, instead of only evaluate the fitness, the multi-objective criterion computes both the 𝒞^*fit*^ score and the proposed 𝒞_*hete*_ score at the same time. Then it updates the *ε*-optimal set and continues the usual splitting routine.

## Results

In this section we compare the proposed MH and MO methods with existing recursive partitioning based heterogeneous treatment effects estimation methods: Regression Tree (RT) [5], Transformed Outcome Tree (TOT) [6], Causal Tree (CT) [7], T-Statistic Tree (TS) [13].

### Synthetic data

The data is generated similarly as previous studies [7, 13]. There are four simulation settings. The first setting focuses on small magnitude of heterogeneities in treatment effects; the second one has higher magnitude than the first; the third one is similar to the second but has significantly more variables; the last one simulates non-linear treatment effects.

Root mean square error (RMSE) and weighted root mean square error (wRMSE) are used for performance evaluation, and the weights for wRMSE are 0.1 if the true and the estimated treatment effects are of the same signs and 1 if otherwise. wRMSE is used because many applications requiring treatment effects estimation are often cost-sensitive. The methods are compared at each split to eliminate the influences from factors such as different stopping criteria. Simulations are repeated 100 times and the average resulted are reported.

#### Algorithm 2: Maintaining an ∈-optimal set

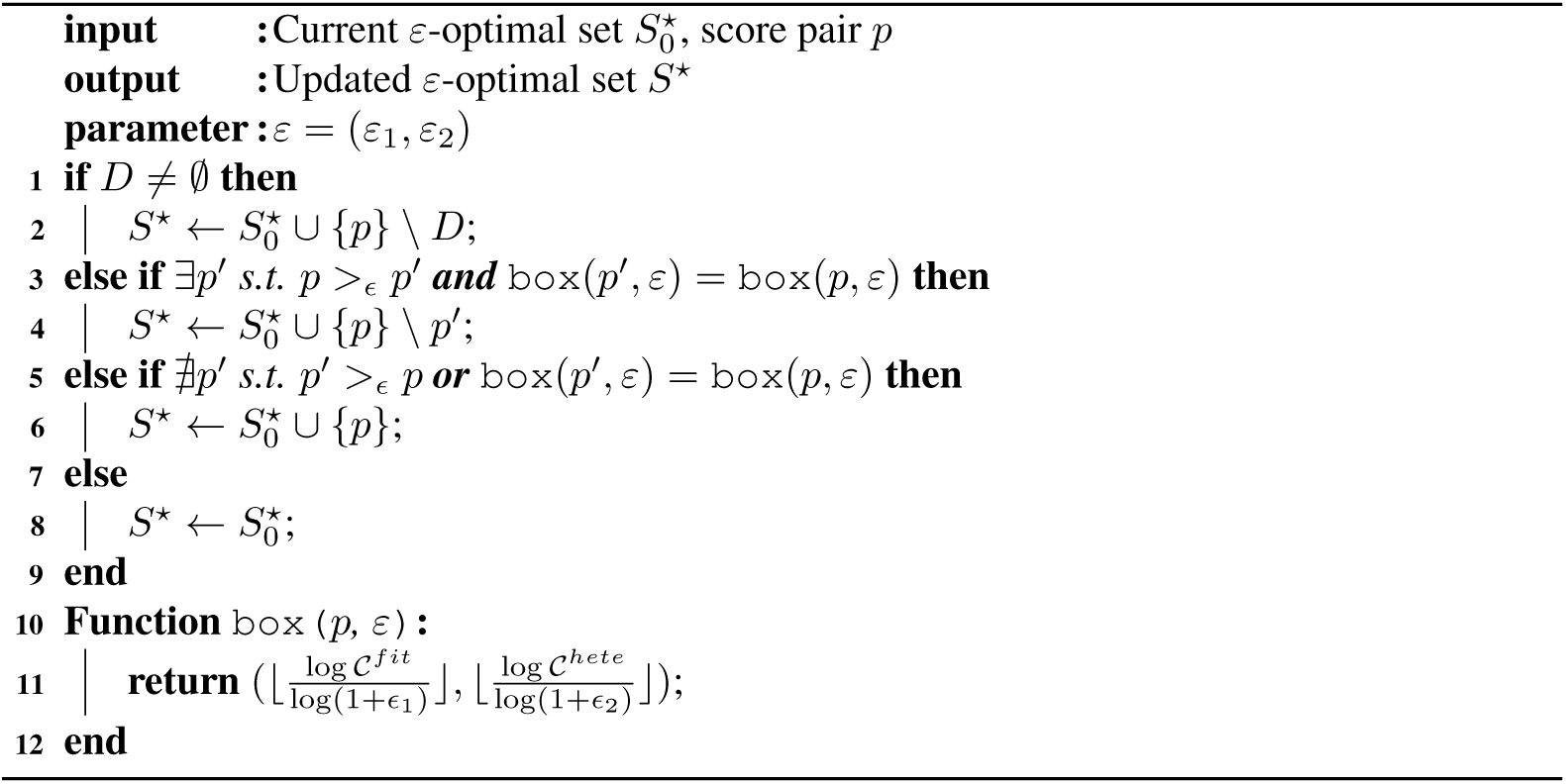

The RMSE and the wRMSE results are shown in Fig. 5 and Fig. 6, respectively. The columns of the figure correspond to different simulation designs, and rows correspond to different samples sizes (*n* = 1000 and *n* = 5000).

**Fig 5.**
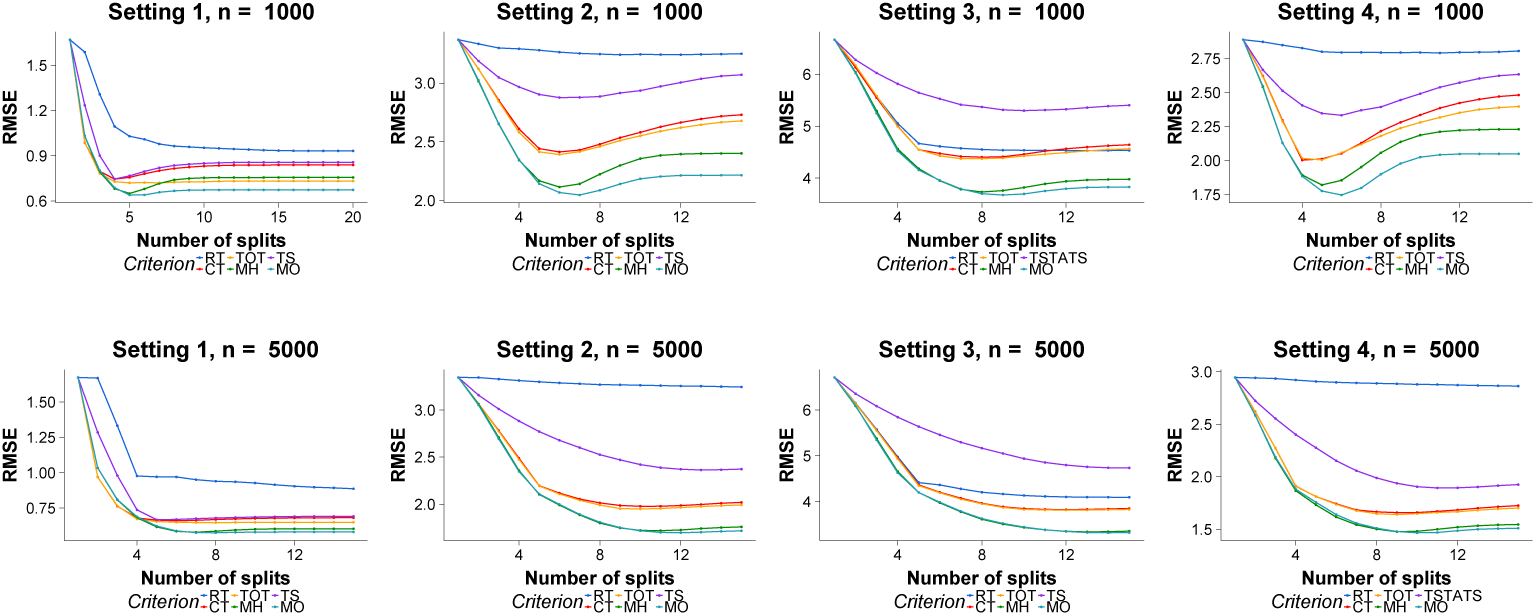
RMSE comparison on simulation datasets. Row: different sample sizes. Column: different experiment designs.

**Fig 6.**
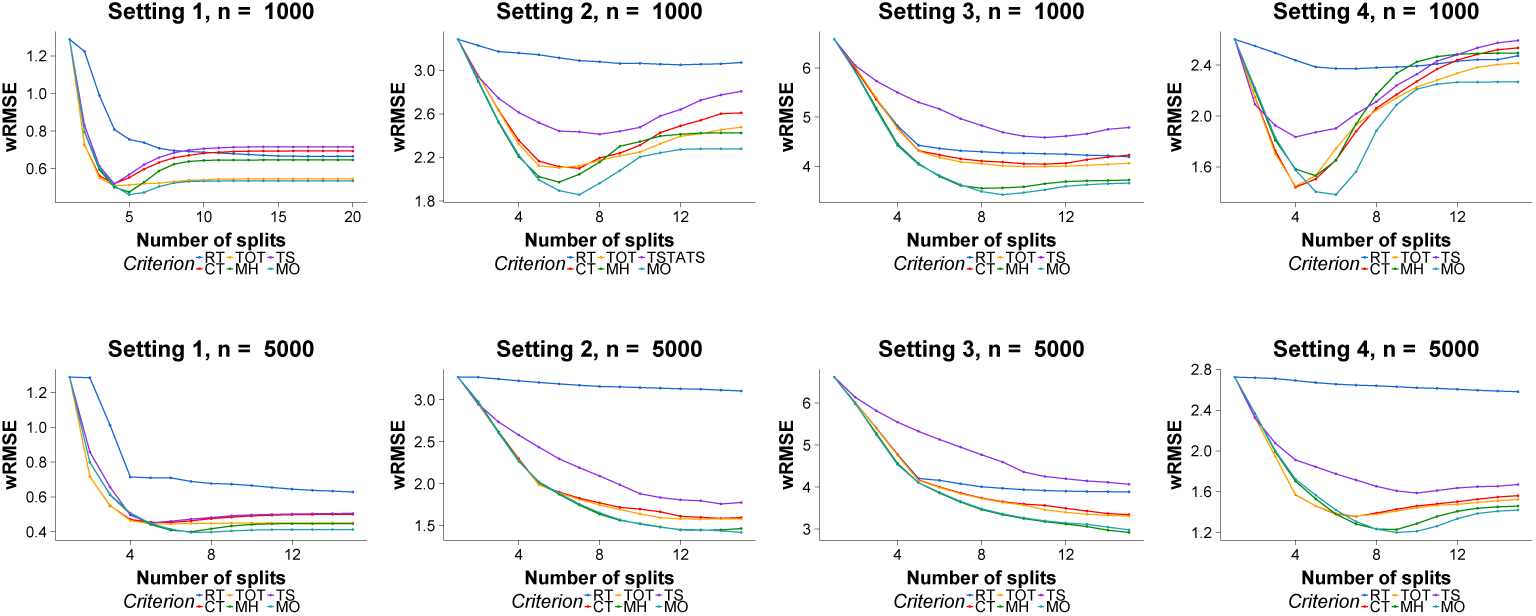
wRMSE comparison on simulation datasets. Row: different sample sizes. Column: different experiment designs.

MH performs better than existing methods when the underlying differences in treatment effects are large. In Setting 2, 3 and 4, MH performs better than all existing methods in terms of RMSE. Although the heterogeneity magnitude is smaller in Setting 1, MH still performs similarly to existing methods. According to the first columns of the figures, the RMSE values of MH is similar to the best existing methods during the first 5 splits. This confirms our previous observation that MH is adept at identifying significant differences in treatment effects because it maximizes heterogeneity explicitly.

The advantage of MH decreases when the underlying heterogeneity magnitude gets smaller. 231 In Setting 1, the simulated treatment effect heterogeneities are much smaller than other settings. 232 As a result, the performances of MH are quite similar to those of existing methods. Furthermore, MH is also vulnerable to spurious heterogeneities. In Setting 1 with *n* = 1000, its performance decrease quickly as the trees grow deeper. When *n* = 1000, both of MH’s measurements are worse than that of TOT after the 6-th splits. When *n* = 5000, although the RMSE values of MH are relatively stable and better than existing methods, the wRMSE values of MH decrease and become identical TOT.

MO performs better than existing methods when the underlying heterogeneities are significantly large. In all four settings, it achieves the lowest RMSE and wRMSE performances in both sample sizes. During the earlier splits, the performances of MO are similar to MH because we prefer score pairs with higher heterogeneity when ties occur.

The advantage of considering heterogeneity and fitness is clearly demonstrated by contrasting MO with MH when the trees grow deeper. After the first few splits, the performances of MH degenerate significantly and the benefit of considering heterogeneity is almost canceled because of over-fitting. In the contrary, the performances of MO continue to improve when MH begins to over-fit the data. When *n* = 100, MO achieves significantly lower error rates than MH in all four simulations settings. Although the differences are closer when the sample size increases, the error rate of MO is still lower than that of MH and existing methods.

In most cases, the performances of existing CATE estimations methods (CT, TS, TOT) are better than the standard regression tree (RT). However, the performances of TS are even worse than RT when the number of variables is large and the sample size is not sufficient, i.e., in Setting 3 when *n* = 1000 and *n* = 5000. This is because TS utilizes statistical tests to decide the split, which suffer from loss of power when the dimensionality grows large. In addition, the situation worsens as the sample size decreases along the tree growth.

It is also worth noticing that in some cases, existing methods have slightly lower wRMSE values than MO. This is related to the strategy of selecting a split point from the *ε*-optimal set. As discussed in the methods section, when there are multiple elements in the *ε*-optimal set, the split with the highest 𝒞^*hete*^ score is chosen from the set. However, if the splitting point with the highest 𝒞^*fit*^ score is selected, the performances of MO will improve in these circumstances, but it will perform worse in other situations. This indicates an adaptive strategy for selecting splitting point from the *ε*-optimal set can further improve the performance.

Different choices of *ε*_1_ and *ε*_2_ values have influence on the performance of MO. Fig. 7 illustrates how the parameter affects the performances in Setting 2 at sample size *n* = 5000. Although the possible values of *ε* range from 0 to 1, in reality only small *ε* values (i.e. [0.05, 0.2]) should should be considered since the parameters proportionally control the amount of tolerance in the *ε*-dominance relationship.

**Fig 7.**
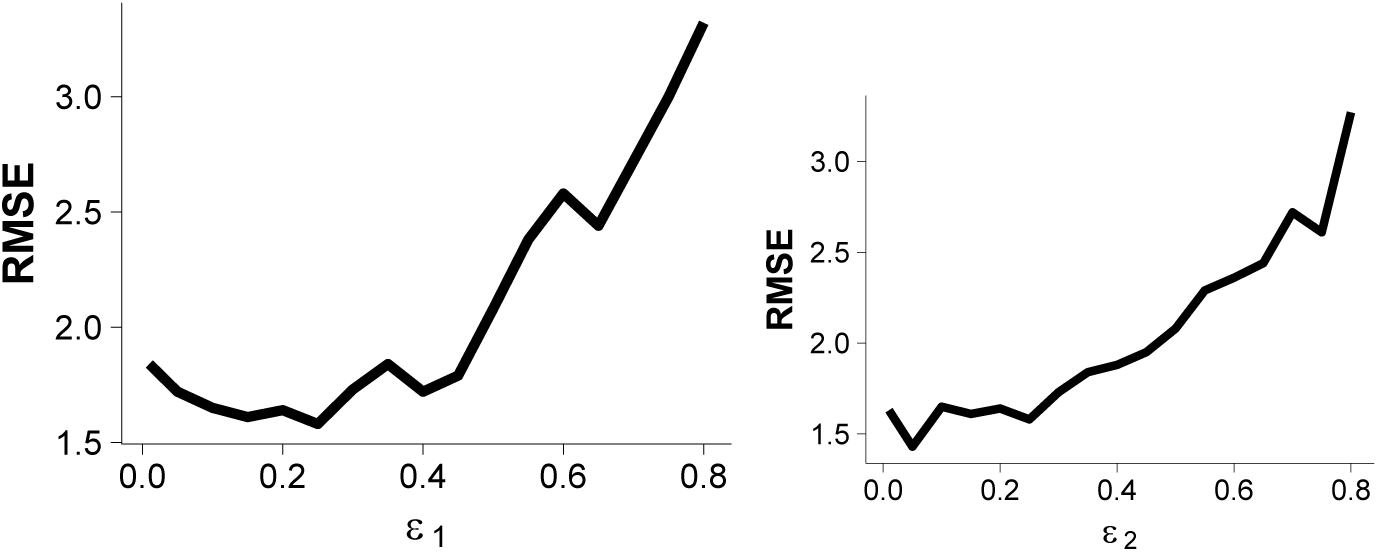
Influence of the parameters ε_1_ and ε_2_.

### Heterogeneous treatment effects of radiotherapy in breast cancer patients

Understanding treatment effect heterogeneity is important to the life quality of cancer patients. More than 50% of the breast cancer patients have received the radiotherapy treatment, equating to over half a million patients worldwide each year. Although radiotherapy is effective for many patients, not all of them benefit from the treatment [14].

In this section we study the heterogeneous treatment effects of radiotherapy on breast cancer patients. The data is obtained from the Cancer Genome Atlas (TCGA) [15]. The radiotherapy status is used as the treatment indicator, the gene expression profiles are used as covariates, and the relapse-free survival status is used as the outcome.

Comparison of CATE estimation algorithms on real-world data is not straightforward because the ground truth treatment effects are not observable and the sample sizes are not large enough to divide the original data into training and testing sets. With this dataset, the performances are evaluated by examining whether the genes selected by each method can be used to differentiate the survival probability between the radiotherapy treated and the untreated patients. An independent test set [16] is used for performance evaluation by examining whether the genes in the models can differentiate the survival probability between the treated and the untreated patients.

In Table I we compare the methods by their *p*-values calculated from log-rank test [17] and the combined *p*-values calculated with the Fisher’s method [18]. Smaller *p*-values indicates that the selected genes are more related to the survival probability of breast cancer patient.

Considering the limited sample size, we restrict the maximum tree size to 4 leaf nodes for each method.

All genes selected by the compared methods are1 related to the heterogeneity of radiotherapy treatment effects. However, as shown in the table, genes selected by MO achieve the smallest *p*-value in all compared methods. It is clear that the genes chosen by MO are clearly the most significantly related to the survival outcomes of breast cancer patients.

An interesting observation is that four out of six methods choose FOXF1 as the first split, indicating that FOXF1 is closely related to breast cancer and the effectiveness of radiotherapy. In biology research, FOXF1 has been recently identified as important cancer-related gene [19]. Our findings could suggest a new direction for exploring its genetic function and contribution in cancer development.

The treatment effect heterogeneities identified by MO are illustrated in Fig. 8, where each panel shows the survival curves comparison between radiotherapy treated and untreated patients. For those patients that are categorized into the first and the second subgroups (first row), their estimated treatment effects of radiotherapy are 0.22 and −0.20, respectively. As evidenced by the *p*-values, the survival probability of the treated is significantly higher than the untreated for patients in the first subgroup, and the survival probability of the treated is significantly lower than the untreated for those in the second subgroup. However, those patients in the third and the last subgroups do not show significant benefits from radiotherapy. Interestingly, according to their negative estimated treatment effects, the prognosis of their disease are likely to worsen following the radiotherapy treatment.

**Fig 8.**
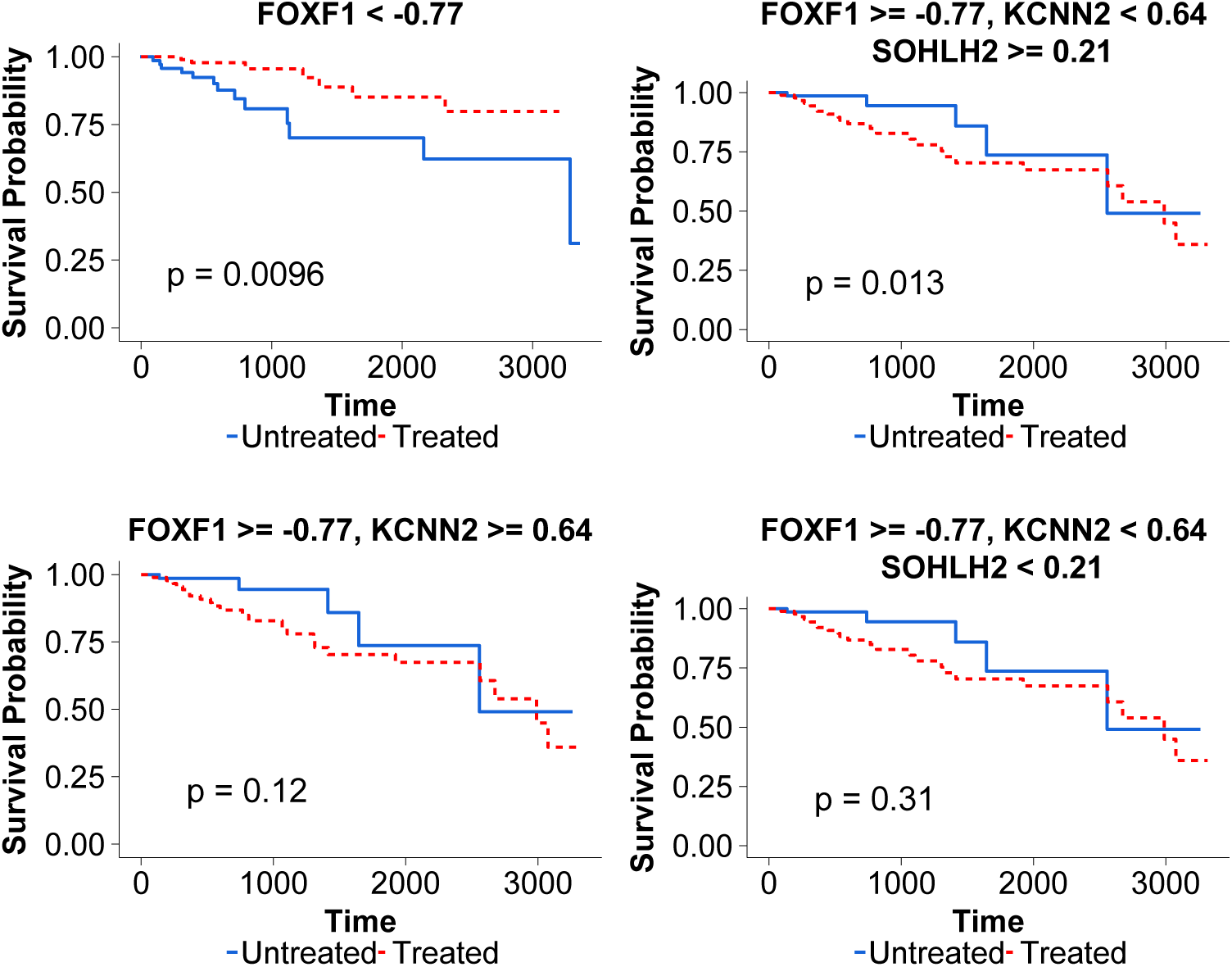
Heterogeneous treatment effects of radiotherapy on breast cancer patients.

**Table 1.** p-values comparison on an independent breast cancer cohort.

| Method | Split 1 | Split 2 | Split 3 | Combined |
| --- | --- | --- | --- | --- |
| RT | 7e-9 | 3e-4 | 0.007 | 7e-12 |
| CT | 1e-16 | 5e-4 | 0.017 | 1e-18 |
| TSTATS | 1e-16 | 5.4e-4 | 0.017 | 1.1e-18 |
| TOT | 6.7e-6 | 2.5e-4 | 0.017 | 9.1e-9 |
| MH | 1e-16 | 1.9e-7 | 0.003 | 9.0e-23 |
| MO | 1e-16 | 1.4e-10 | **0.001** | **3.5e-26** |

## Related works

**RT** [5]. Standard regression tree can be modified to estimate heterogeneous treatment effects [7]. 310 Specifically, the tree is constructed using the CART algorithm, and the treatment effect 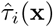 is estimated according to Equation 2 using the samples within the same leaf.

**Transformed Outcome Tree** [6]. Transformed Outcome Tree (TOT) is based on the insight that existing regression tree methods can be used to estimate treatment effect by utilizing a transformed version of the outcome variable 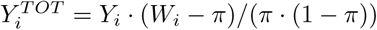 as the regression target. Because 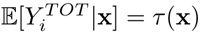, standard regression tree can be applied to the transformed outcome where the estimation of the sample average of 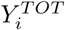 within each leaf can be interpreted as the estimation of the treatment effects.

**Causal Tree** [7]. Causal Tree (CT) seeks the splitting point using the fitness criterion, but it does not consider the heterogeneity. In addition, they propose to divide the training samples into two disjoint parts to avoid bias in the treatment effect estimation, where the first part is used for selecting split and the second part is used to estimate the treatment effects in the model.

**Squared t-Statistic Tree** [13]. squared T-Statistic tree (TS) seeks the split with the largest value for the square of the t-statistic for testing the null hypothesis that the average treatment effect is the same in the two potential leaves. The criterion is defined as:

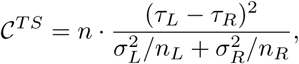

where *σ*^2^ is the variance of treated and untreated samples within a node.

𝒞**^*hete*^** criterion is different from the TS criterion. Because the sample size grows smaller as the tree grows, the statistical test used in [13] suffers from loss of power. Unless the subgroup treatment effects are quite large, this method often fails to detect the effects in subgroups [20]. In the experiments it is demonstrated that the performance of TS degenerates significantly as the number of variables increases, whereas the performances of MH remain unaffected.

## Conclusion

Previous studies of heterogeneous treatment effect estimation focus on improving the fitness of the model, which may lead to less informative models. In this paper, we demonstrate that heterogeneity needs to be considered explicitly. Moreover, heterogeneity and fitness are needed simultaneously in order to achieve a reliable and informative model.

MH performs well when the underlying heterogeneities are large and MO performs consistently better than existing ones. However, there is still much to explore. Particularly, an adaptive strategy for choosing the most suitable elements from the *ε*-optimal set could further improve the performances.

## Acknowledgments

This work is supported by Australian Research Council Discovery Project (DP140103617). Thuc Duy Le is supported by NHMRC Grant (ID 1123042).

